# Essential role of astrocytes in rapid activity-dependent restructuration of the axon initial segment

**DOI:** 10.1101/2025.07.31.667937

**Authors:** Rafael Sanz-Gálvez, Yanis Inglebert, Arlette Kolta

**Author notes:** These authors contributed equally to this work and share first authorship (Rafael Sanz-Gálvez, Yanis Inglebert).

## Abstract

One key regulator of neuronal excitability is the axon initial segment (AIS), a highly specialized axonal region, enriched in ion channels, where action potentials are initiated. The AIS can undergo significant morphological changes to fine-tune neuronal excitability in response to external perturbations. Long considered solely a homeostatic mechanism operating over long timescales (hours to days) to adjust excitability, we show here that this phenomenon can also occur rapidly, within minutes, following a brief period of high activity in layer 5 pyramidal neurons of the visual cortex. Because astrocytes have been known to regulate neuronal excitability, we explored the effects of gliotransmitters on this process and identified the calcium-binding protein S100β from astrocytes to be required for the rapid reorganization of the AIS.

Graphical abstract

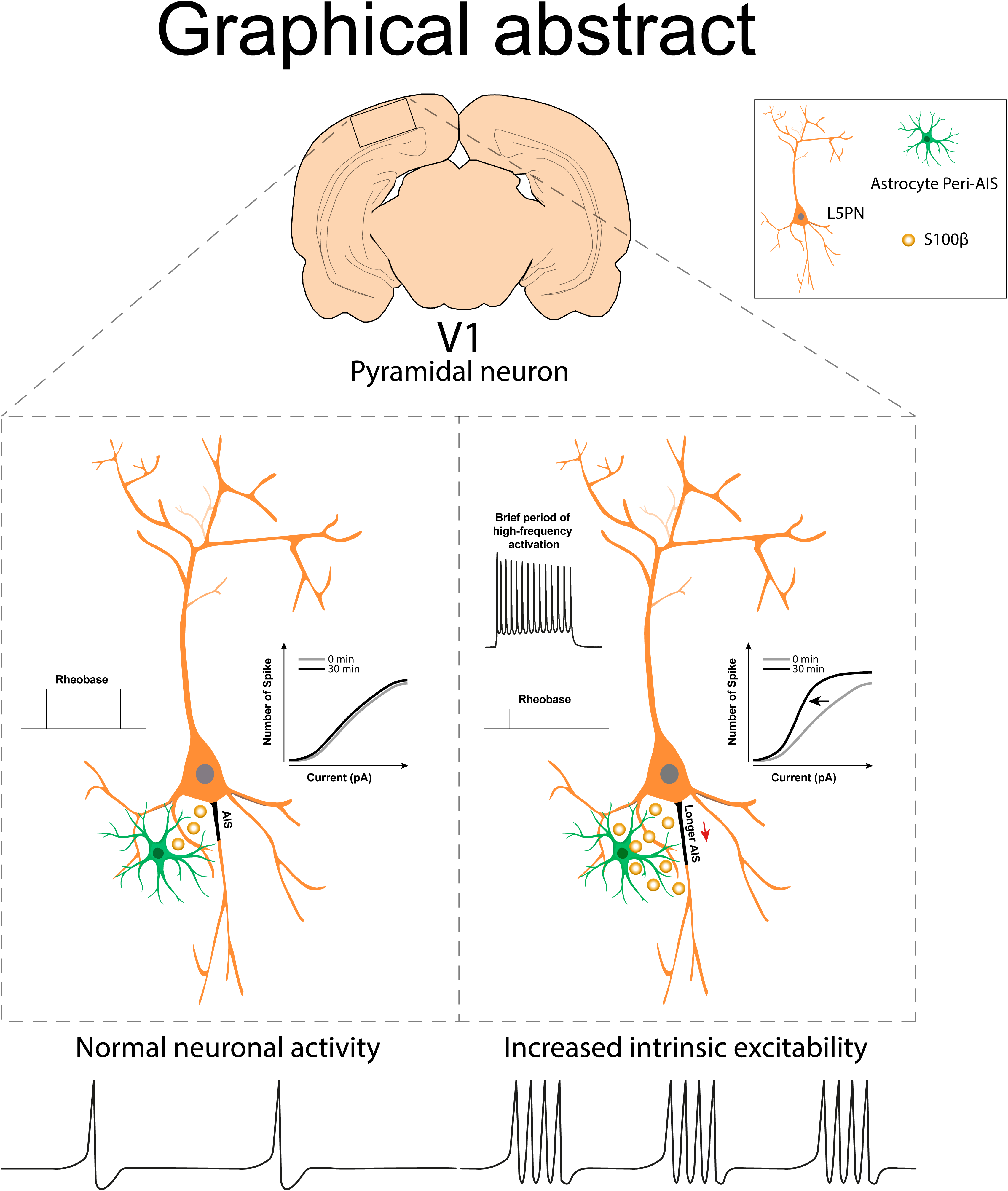

## INTRODUCTION

The plasticity of the input-output function is regulated by diverse plasticity rules to adapt neuron’ excitability in response to external or internal perturbations. One powerful regulator is the axon initial segment (AIS), a specialized subcellular region of the proximal axon where action potentials are shaped and initiated (Debanne et al., 2011). Structural plasticity of the AIS can influence neuronal output as demonstrated by changes in AIS length in response to external perturbation to maintain appropriate neuronal excitability (Yamada and Kuba, 2016; Fréal et al., 2023). The length of the AIS is a reliable indicator of neuronal excitability, where increased length enhances excitability and decreased length lowers it. Plasticity of the AIS is highly used in different sensory processing brain regions to fine-tune neuronal excitability or during development to maintain stable network excitability (Kuba et al., 2006; Schlüter et al., 2017). For instance, in the visual cortex, the AIS is developmentally regulated by visual experience during critical period (Gutzmann et al., 2014; Höfflin et al., 2017). Historically, AIS plasticity has been considered a homeostatic mechanism operating over long timescales (days to weeks), allowing neurons to maintain their firing set point in response to network perturbations. This view largely stems from studies conducted in neuronal cultures following chronic network manipulations (Grubb and Burrone, 2010). This perspective has evolved over time, with plasticity being described at progressively shorter timescales, from hours to minutes (Grubb et al., 2011; Evans et al., 2015). The latest findings, using newly developed genetic tools (Thome et al., 2025) that label the scaffolding protein Ankyrin G (AnkG), a specific marker of the AIS (Leterrier, 2018) reveal that AIS reorganization can occur within minutes following long-term depression in the hippocampus (Fréal et al., 2023). This is the first evidence showing that the synergistic excitability changes observed following synaptic plasticity could be accompanied by AIS plasticity (Campanac and Debanne, 2007, 2008). However, synaptic plasticity is not the sole mechanism of memory storage; intrinsic excitability plasticity can also occur in response to non-homeostatic stimulation and could be associated as well with rapid AIS plasticity, a possibility that, until now, has never been tested. Here, we provide initial evidence of a close relationship between intrinsic excitability changes and AIS plasticity. In Layer 5 cortical pyramidal neurons (L5PNs), repetitive bursts high-frequency firing induce long-term potentiation of intrinsic excitability (LTP-IE) a fundamental mechanism to refine visual circuit (Cudmore and Turrigiano, 2004; Nataraj et al., 2010; Duménieu et al., 2025). In cortical circuit, LTP-IE is dependent on extracellular calcium influx and modulation of ion channels (Cudmore and Turrigiano, 2004). Among the various external factors that could influence LTP-IE, astrocytes have long been overlooked despite increasing evidence of their role as powerful modulators of input-output functions and sensory processing (Sanz-Gálvez et al., 2024). Recent findings show that astrocytes directly interact with the axon or the AIS to modulate excitability through the release of gliotransmitters (Debanne and Rama, 2011; Lezmy et al., 2021). Moreover, in murine models of Alzheimer’s disease, AIS integrity and length deteriorate due to transcriptomic and proteomic alterations in astrocytes (Benitez et al., 2024). One relatively understudied gliotransmitter, S100β, is particularly intriguing as it functions both as a calcium (Ca²⁺)-binding protein to regulate the concentration of extracellular calcium and a modulator of various voltage-gated ion channels (Hermann et al., 2012; Bancroft et al., 2022). Extracellularly, it primarily binds to membrane receptors such as the multi-ligand Receptor for Advanced Glycation End Products (RAGE), triggering intracellular signaling cascades that, depending on its concentration, may lead to physiological or pathological outcomes (Donato et al., 2009). We have previously shown that S100β and stimulation of astrocytes (presumably through release of S100β) elicit spiking in L5PNs from the visual cortex and trigeminal primary afferent axons or promote rhythmic activity in trigeminal sensori-motor circuit through the enhanced activation of voltage-gated sodium (NaV) 1.6 channels (Morquette et al., 2015; Ryczko et al., 2021; Gaudel et al., 2025). Interestingly, NaV 1.6 is one of the main isoforms of NaV channels expressed at the AIS in excitatory neurons which distribution is reorganized during plasticity (Fréal et al., 2023). Here, we first test whether LTP-IE is associated with rapid AIS plasticity in L5PNs of the visual cortex and then examine if astrocytes regulate this process through the release of the gliotransmitter S100β.

## RESULTS

### Long-lasting potentiation of intrinsic excitability (LTP-IE) is associated with rapid AIS plasticity

Layer 5 pyramidal neurons (L5PNs) were recorded in acute slices from the primary visual cortex (V1) obtained from Vglut2-AnkG-GFP young mice (P14–P30) (see Methods and **Fig. 1A**), and their intrinsic excitability was monitored by a 500-ms constant direct current pulse every 10 seconds for 5 min after which repetitive current injection (500-ms depolarizing pulses, 60 times at 4-second interval, **Fig. 1B bottom)** was used to induce long-lasting potentiation of intrinsic excitability (LTP-IE). All experiments were conducted in the presence of glutamatergic and GABAergic receptors blockers to prevent interferences from synaptically released fast excitatory and inhibitory neurotransmitters. As previously described (Cudmore and Turrigiano, 2004), this protocol induced a robust and long-lasting increase in intrinsic excitability (151% ± 12, n=6, **Fig. 1B**), which was monitored every 10 seconds for 30 minutes after induction. After confirming our immunohistochemical results showing that the GFP signal colocalized with the master AIS protein, AnkG (**Fig. 1A top**), we proceeded to monitor AIS remodeling in real-time before and after the induction protocol in the recorded neuron filled with Alexa Fluor 647 (**Fig. 1A bottom**). Fluorescence images of AnkG-GFP taken 30 minutes after induction revealed a rapid and significant elongation in the distal part of the AIS, as confirmed by fluorescence intensity measurements (mean AIS length: 29.68 ± 1.25 µm before induction and 33.29 ± 1.35 µm after induction, n=6, t-test, P<0.001; **Fig. 1C**). To avoid rapid deterioration due to phototoxicity, imaging of the AIS was minimized, but in a separate set of experiments (n=5), images were captured every 5 minutes, revealing that in some cells, the increase began as early as 10–15 minutes after induction **(Fig. S1)**. In the absence of the induction protocol, no LTP-IE (102% ± 2, n=5, **Fig. 1B**) or AIS remodeling was observed (mean AIS length: 34.31 ± 0.95 µm before and 34.21 ± 0.92 µm after, n=5, t-test, P>0.05; **Fig. 1F**). To determine changes in neuronal excitability, the number of APs for each increase in injected current was counted, establishing input-output curves before and after the induction protocol. Additionally, rheobase, an important parameter of neuronal excitability, was measured. In neurons subjected to the brief repetitive firing period, a significant increase in intrinsic excitability was observed across other supra-threshold current amplitudes (from 30 pA to 100 pA, t-test, P<0.01; P<0.05; **Fig. 1D**), whereas no significant changes were observed in control neurons (all steps; t-test, P>0.05; **Fig. 1G**). Moreover, in neurons subjected to the induction protocol, but not in control neurons (**Fig. 1H**), a significant decrease in rheobase was observed (30 ± 12 pA before vs. 21 ± 9 pA after induction, n=6, t-test, P<0.01; **Fig. 1E**). Together, these data demonstrate that a period of high-frequency activation as brief as 4 min suffice to induce remodeling of the AIS and LTP-IE in L5PNs.

**Figure 1.**
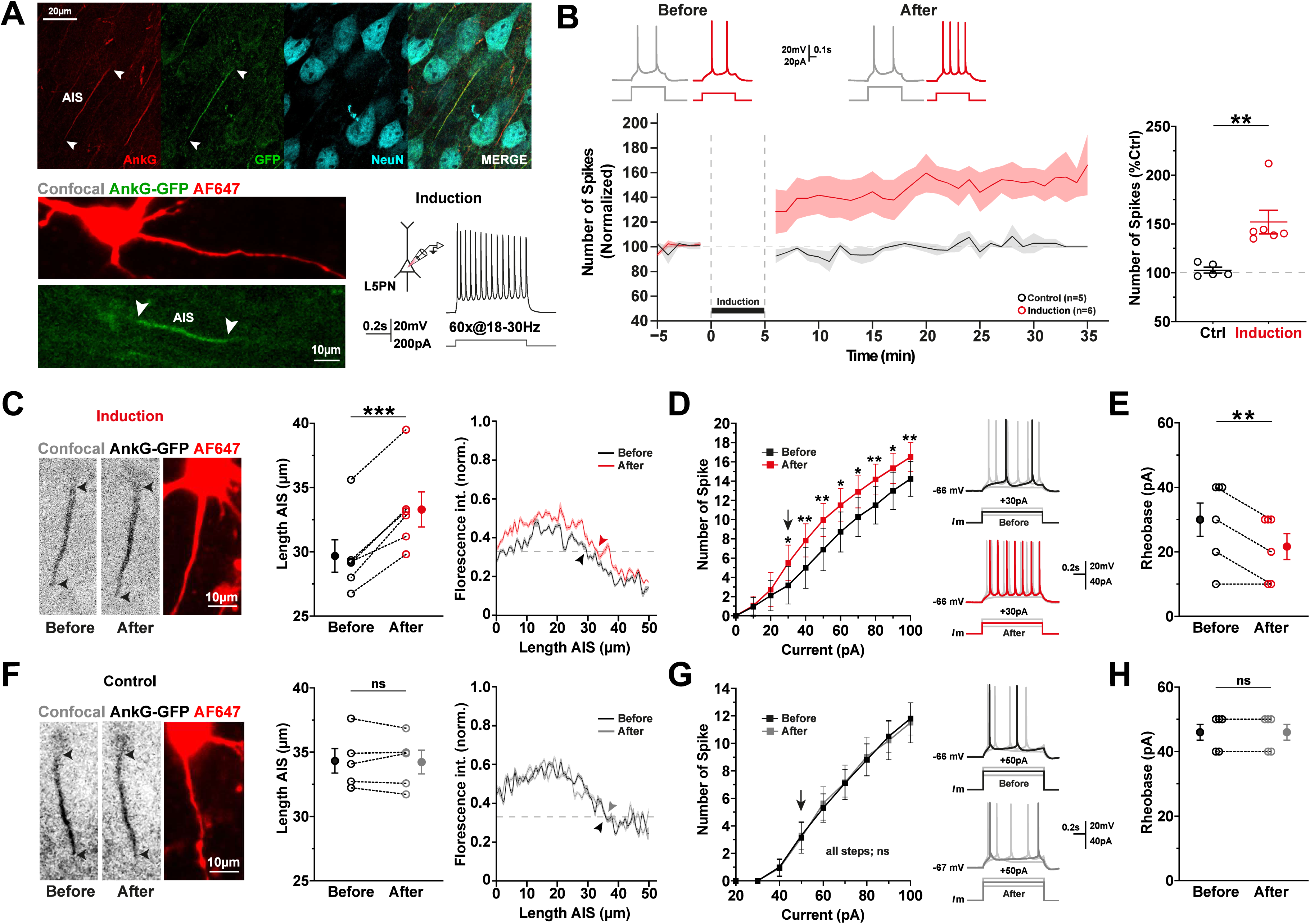
LTP-IE is associated with rapid AIS plasticity. **A.** *Top*. High-resolution images showing strong co-localization of AnkG (red, *left*) and GFP (green, *middle*) in pyramidal neurons (cyan, *right*) from layer 5 (L5PNs) of the visual cortex. *Bottom left.* Photomicrography of the axon initial segment (AIS) (AnkG-GFP) in L5PNs filled with Alexa 647. *Bottom right.* Membrane response of the whole-cell recorded neuron at the soma during the induction protocol. **B.** *Left*. Time course of normalized spike number before and after (representative traces shown above) induction (red) and in control conditions (black). *Right*. Quantification of spike number changes in the last 10 minutes, with individual values for each cell represented as red (induction) and black (control) open circles. **C.** *Left*. Live image of an AnkG-GFP-positive neuron filled with Alexa Fluor 647 before and after induction. Black arrowheads indicate the start and end of the AIS. *Middle*. Induction of intrinsic excitability potentiation elongates AIS length (left, manual measurement of individual cells; right, mean normalized fluorescence intensity profiles before (black) and 30 min after induction (red). Arrowheads indicate AIS end positions where the fluorescence profile crosses the 33% threshold, showing AIS elongation). **D.** *Left.* Current-number of spike relationships for each condition (before and after induction). Induction caused a leftward shift in the mean curve. *Right.* Representative traces of individual current injections at different amplitudes. Note the difference in the spike number with the same amplitude before and after induction. **E.** Decrease in Rheobase after induction. **F.** *Left*. Live image of an AnkG-GFP-positive neuron fixed with Alexa Fluor 647 before and after the control condition. Black arrowheads indicate the start and end of the AIS. *Middle*. No AIS length change conditions (left: manual measurements; right: fluorescence profiles before and after control, with no shift at the 33% threshold). **G.** *Left.* Current-number of spike relationships for each condition (before and after the control period). No shift in the mean curve was observed. *Right.* Representative traces of individual current injections at different amplitudes in the control condition, with no changes observed. **H.** No change in Rheobase in the control condition.

### LTP-IE and AIS plasticity involves peri-AIS astrocyte signaling

Astrocytes are involved in various forms of synaptic plasticity (Yang et al., 2003; Andersson et al., 2007; Falcón-Moya et al., 2020; Sanz-Gálvez et al., 2024); however, their potential contribution to intrinsic plasticity has not been assessed. Therefore, we examined whether peri-AIS astrocytes (astrocytes near the AIS) are necessary for LTP-IE and the consequent structural remodeling of the AIS in L5PN neurons. To address this, we preincubated acute brain slices for 5 minutes RT with Sulforhodamine (SR101, 1 µM), a widely used astrocyte marker in different brain regions (**Fig. 2A**) (Watanabe et al., 2023). Next, a single astrocyte was loaded with 20 mM of the Ca²⁺ chelator BAPTA via a patch pipette to inhibit Ca^2+^-dependent vesicular release and gliotransmitter signaling (Araque et al., 1998) and a nearby L5PN (distance between the AIS of the recorded neuron and the patched astrocyte; 37.02 ± 8.74 µm, **Fig. 2B**) was targeted for whole-cell recording after allowing time (15 min) for BAPTA to diffuse in the astrocytic syncitium. Astrocytic inactivation with BAPTA prevented the induction of LTP-IE (103% ± 4, n=5; **Fig. 2B**) and the associated AIS lengthening (mean AIS length: 33.4 ± 0.95 µm before induction vs. 32.61 ± 0.94 µm after induction, n=5, t-test, P>0.05; **Fig. 2C**). No significant changes were observed in input-output curves or rheobase values before and after induction (**Fig. 2D,E**). Taken together, these results clearly indicate that peri-AIS astrocytes are required for the induction of this form of intrinsic excitability plasticity and AIS remodeling.

**Figure 2.**
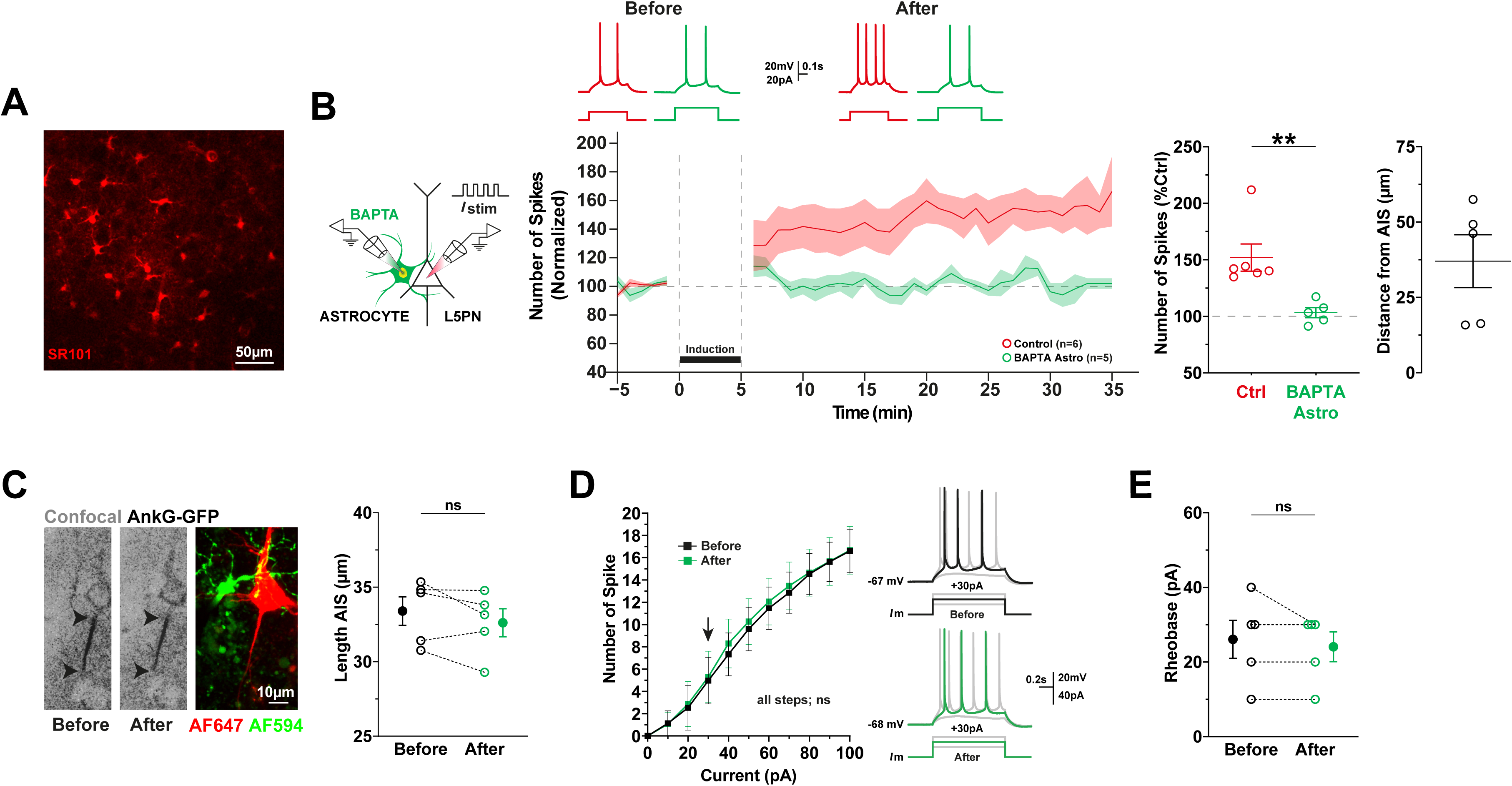
Astrocytic inactivation prevents LTP-IE and AIS plasticity. **A.** Confocal imaging of Sulforhodamine (SR101) labeling in astrocytes of layer 5 of the visual cortex. **B.** *Left.* Experimental paradigm. Stimulation was performed by current injection into the whole-cell recorded neuron, after targeting a neighboring peri-AIS astrocyte for recording with a BAPTA filled pipette for its inactivation. *Middle.* Time course of normalized spike number before and after induction (red) and during peri-AIS astrocyte network inactivation (green), with representative traces shown above. *Right.* Quantification of spike number changes in the last 10 minutes, with individual values for each cell represented as red (induction) and green (astrocytes inactivated) open circles and distance of the inactivated astrocytes from the soma of the recorded neuron. **C.** Live image of an AnkG-GFP-positive neuron filled with Alexa Fluor 647 before and after induction during peri-AIS astrocyte network inactivation, filled with Alexa Fluor 594. Black arrowheads indicate the start and end of the AIS. *Right.* Peri-AIS astrocyte inactivation prevented the change in AIS length. **D.** *Left.* Current-number of spike relationships for each condition (before and after induction during peri-AIS astrocyte network inactivation). No shift in the mean curve was observed. *Right.* Representative traces of individual current injections at different amplitudes, with no changes observed. **E.** No change in Rheobase in the recorded neuron during peri-AIS astrocyte network inactivation.

### LTP-IE and AIS plasticity depend on the astrocytic protein S100β

In the visual cortex, LTP-IE, as other types of synaptic plasticity, depends on extracellular Ca^2+^ (Inglebert et al., 2020), since it is abolished in 0mM Ca^2+^ medium (Cudmore and Turrigiano, 2004). Our previous studies demonstrated that astrocytes could tightly regulate extracellular Ca^2+^ levels through the release of S100β (Morquette et al., 2015) and that optogenetic activation of layer 5 astrocytes in the visual cortex decreases extracellular Ca^2+^ via S100β release (Ryczko et al., 2021). Based on these findings, we investigated whether this Ca^2+^-binding astrocytic protein could locally modulate LTP-IE and AIS structural remodeling. To this end, we locally administered an anti-S100β antibody in the vicinity of the AIS for 5 minutes prior to induction to prevent the endogenous protein from exerting its effects and from binding Ca^2+^ (Morquette et al., 2015). Applications of the same antibody, but denaturated by preheating to 100 degrees Celsius for 20 minutes, served as controls. The S100β antibody effectively blocked LTP-IE, whereas the denatured antibody did not preclude its induction (S100β antibody: 106% ± 17, n=5 vs. S100β antibody denatured: 180% ± 9, n=6, (MW) U test, P<0.05; **Fig. 3A**). Confocal imaging further revealed that the S100β antibody prevented AIS elongation (mean AIS length: 34.76 ± 0.41 µm before induction and 34.48 ± 0.48 µm after induction, n=5, t-test, P>0.05; **Fig. 3B**). Conversely, with the denatured antibody, significant AIS elongation was observed (mean AIS length: 32.45 ± 1.67 µm before induction and 34.63 ± 1.62 µm after induction, n=6, t-test, P<0.001; **Fig. 3E**) similar to what is observed in control condition (**Fig 1C**). The S100β antibody application was found to cause a small decrease of excitability as measured in the input-output curves, but this change was not statistically significant (all steps; t-test, P>0.05; **Fig. 3C**). This effect is likely attributable to an increase in extracellular Ca^2+^ levels induced by S100β blockade, which is consistent with prior findings indicating that elevated extracellular Ca^2+^ levels suppress neuronal excitability (Forsberg et al., 2019, 2025). In contrast, the denatured antibody application is associated with an upward shift of the input-output curve (pA40, pA60-pA100, t-test, P<0.05; **Fig. 3F**). No significant changes were observed in rheobase under any condition (**Fig. 3D,G**). Collectively, these data underscore the necessity of the Ca^2+^-binding astrocytic protein S100β for the induction of LTP-IE and AIS morphological modifications in LPN5s.

**Figure 3.**
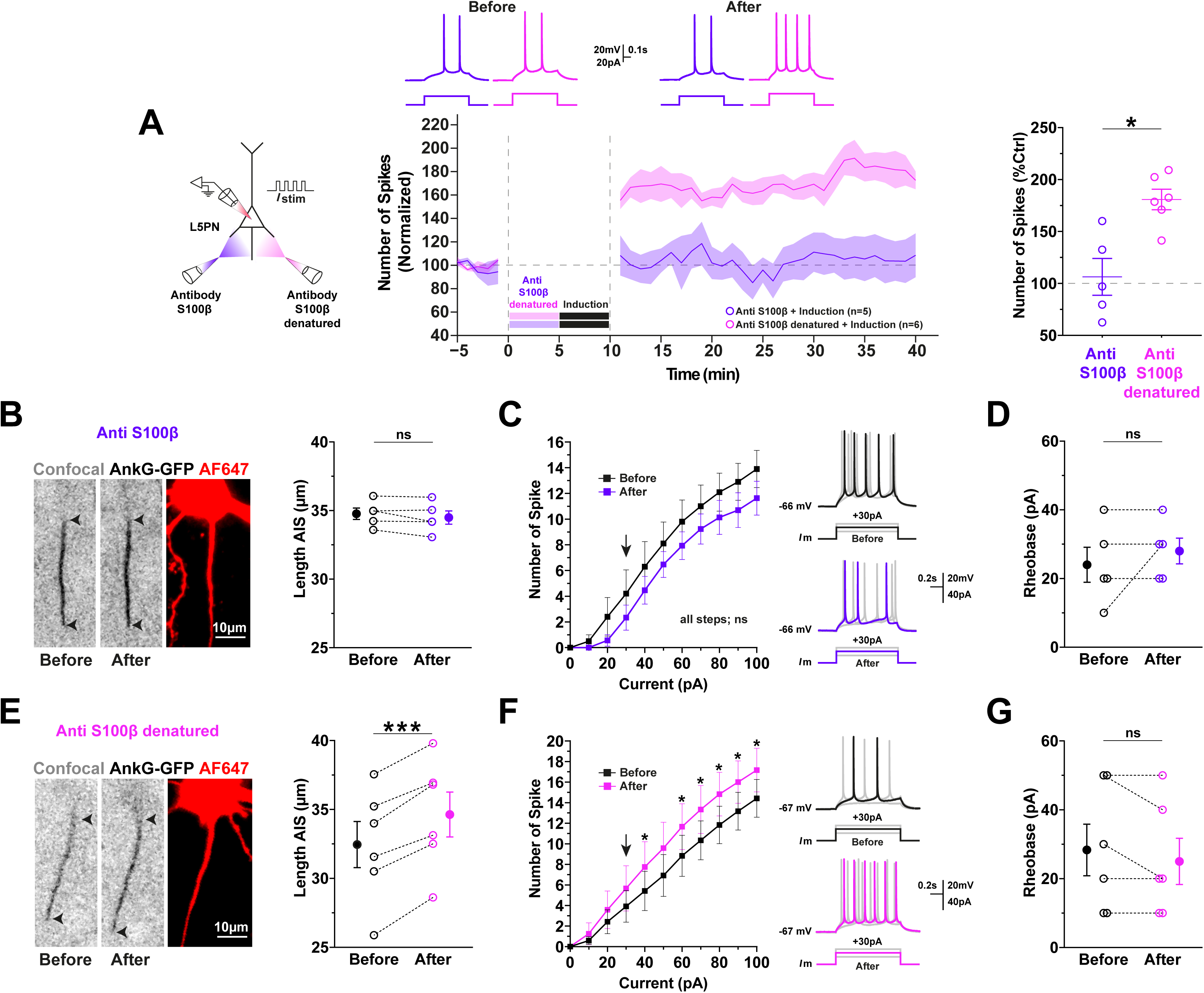
Blockade of the astrocytic protein S100β impairs LTP-IE and AIS plasticity. **A.** *Left.* Experimental paradigm. The induction protocol was performed by current injection into the whole-cell recorded neuron 5 minutes after a local application of either an S100β antibody or its denatured form. *Middle.* Time course of normalized spike number before and after induction, with prior application of S100β antibody (purple) and its denatured form (pink), with representative traces shown above. *Right.* Quantification of spike number changes in the last 10 minutes, with individual values for each cell represented as purple (S100β antibody) and pink (S100β antibody denatured) open circles. **B.** *Left*. Live image of an AnkG-GFP-positive neuron filled with Alexa Fluor 647 before and after the application of S100β antibody, followed by the induction protocol. Black arrowheads indicate the start and end of the AIS. *Right.* The application of the S100β antibody, prior to induction, prevents the change in AIS length. **C.** *Left.* Current-number of spike relationships for each condition (before and after S100β antibody application, followed by the induction protocol). Induction caused a rightward shift in the mean curve. *Right.* Representative traces of individual current injections at different amplitudes. Note the difference in the spike number with the same amplitude before and after induction with prior application of S100β antibody. **D.** No change in Rheobase in the recorded neuron with the application of S100β antibody prior to induction. **E.** *Left*. Live image of an AnkG-GFP-positive neuron filled with Alexa Fluor 647 before and after the application of S100β antibody denatured, followed by the induction protocol. *Right.* The application of denatured S100β antibody prior to induction does not prevent the change in AIS length. **F.** *Left.* Current-number of spike relationships for each condition (before and after S100β antibody denatured application, followed by the induction protocol). Induction caused a leftward shift in the mean curve. *Right.* Representative traces of individual current injections at different amplitudes. Note the difference in the spike number with the same amplitude before and after induction with prior application of S100β antibody denatured. **G.** No change in Rheobase in the recorded neuron with the application of S100β antibody denatured prior to induction.

### Optogenetic activation of peri-AIS astrocytes induces LTP-IE and AIS plasticity through S100β release

Having established that astrocytes and S100β are critically involved in mediating LTP-IE and AIS remodeling, we next aimed to determine whether selective activation of peri-AIS astrocytes alone could suffice to induce LTP-IE. To address this, neonatal VGlut2-AnkG-GFP mice (P0) were bilaterally injected intracerebrally with an adeno-associated virus (AAV) expressing Channelrhodopsin-2 (ChR2) fused to the mCherry reporter and targeted to astrocytes using the GFAP promoter (**Fig. 4A left**). To verify that viral transfection was specifically restricted to astrocytes, immunohistochemistry was performed on V1 sections using an antibody against S100β. Immunostaining revealed that viral expression was targeted to astrocytes, as evidenced by the colocalization of mCherry/ChR2-GFAP with S100β (**Fig. 4A right**). Next, in acute brain slices, patch-clamp recordings demonstrated that transfected astrocytes responded to light pulses (using a combination of 440 nm and 488 nm lasers from a confocal microscope) by evoking membrane depolarizations (**Fig. S2**). This response was verified in representative experiments (**see Methods**). Additionally, we confirmed that viral transfection did not affect the real-time visualization of AIS morphology (**Fig. 4B bottom left**).

**Figure 4.**
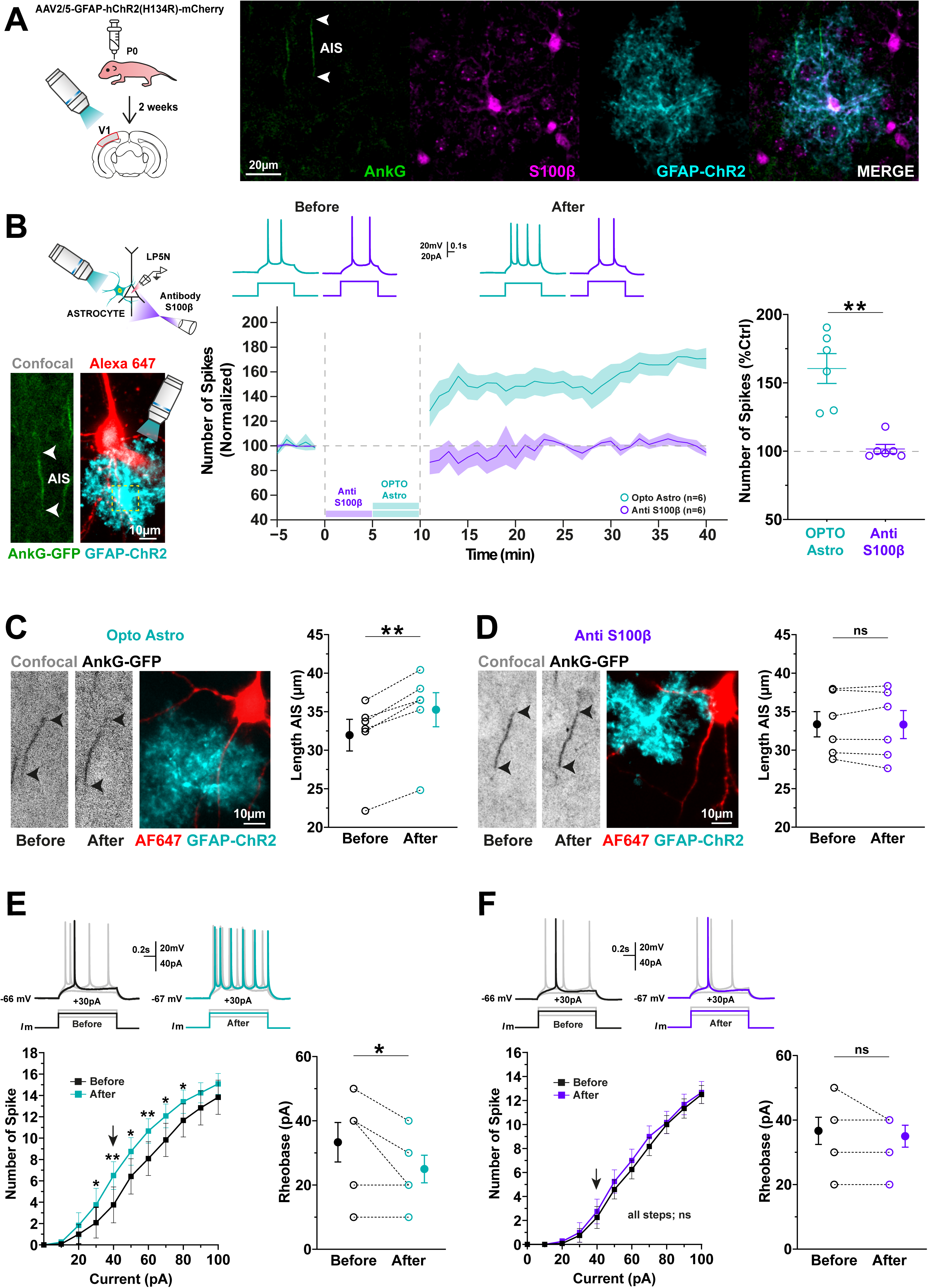
Optogenetic activation of peri-AIS astrocyte reproduces LTP-IE and AIS plasticity. **A.** *Left*. Schematic representation of the strategy used for the optogenetic activation of astrocytes in layer 5 of the cortex visual. *Right*. Confocal imaging of immunohistochemistry-confirmed AAV GFAP-ChR2 (cyan) expression in astrocytes (S100β, magenta) in AnkG-GFP-positive mice (green). **B.** *Left*. Photomicrography of the AIS (AnkG-GFP) in an L5PN filled with Alexa 647 alongside an astrocyte expressing AAV GFAP-ChR2. **B.** *Middle*. Time course of normalized spike number before and after optogenetic activation of astrocytes, with (purple) or without (cyan) S100β antibody application, with representative traces above. *Right.* Quantification of spike number changes in the last 10 minutes, with individual values for each cell represented as cyan (optogenetic activation of astrocytes), purple (S100β antibody followed by optogenetic activation of astrocytes). **C.** *Left*. Live image of an AnkG-GFP-positive neuron filled with Alexa Fluor 647 before and after optogenetic activation of an astrocyte expressing ChR2 (cyan). Black arrowheads indicate the start and end of the AIS. *Right.* The optogenetic activation of astrocytes significantly elongates the length of the AIS. **D.** *Left.* Live image of an AnkG-GFP-positive neuron fixed with Alexa Fluor 647 before and after optogenetic activation of a ChR2-expressing astrocyte (cyan), with prior S100β antibody application. Black arrowheads indicate the start and end of the AIS. *Right.* The application of the S100β antibody, prior to optogenetic peri-AIS astrocyte activation, prevents changes in AIS length. **E.** *Left.* Current-number of spike relationships for each condition (before and after optogenetic activation of astrocytes). Optogenetic activation caused a leftward shift in the mean curve. *Top.* Representative traces of individual current injections at different amplitudes. Note the difference in the spike number with the same amplitude before and after optogenetic activation of astrocytes. *Right*. Change in Rheobase in the recorded neuron during optogenetic peri-AIS astrocyte activation. **F.** *Left.* Current-number of spike relationships for each condition (before and after optogenetic astrocyte activation with prior S100β antibody application). *Top.* Representative traces of individual current injections at different amplitudes, with no changes observed. *Right*. No change in Rheobase in the recorded neuron during optogenetic peri-AIS astrocyte activation with prior S100β antibody application.

Subsequently, excitability of L5PNs was assessed as above, but the induction protocol was substituted by photostimulation of astrocytes (short light pulses of 2 seconds on, followed by 2 seconds off, repeated 60 times; **Fig. S2**) which was as efficient for induction of LTP-IE (**Fig. 4B**). Importantly, local administration of the S100β antibody for 5 minutes prior to optogenetic stimulation of peri-AIS astrocytes abolished LTP-IE (opto Astro: 160% ± 10, n=6 vs. S100β antibody + opto Astro: 101% ± 3, n=6; (MW) U test, P<0.01; **Fig. 4B**). Confocal imaging revealed a significant elongation of the AIS length 30 minutes after optogenetic stimulation of peri-AIS astrocytes (mean AIS length: 31.96 ± 2.05 µm pre-stimulation vs. 35.25 ± 2.20 µm post-stimulation, n=6, t-test, P<0.01; **Fig. 4C**), which was prevented when the S100β antibody was applied locally (mean AIS length: 33.14 ± 1.98 µm pre-stimulation vs. 32.85 ± 2.15 µm post-stimulation, n=6, t-test, P>0.05; **Fig. 4D**). Input-output curves showed that responses to suprathreshold current amplitudes significantly increased after optogenetic stimulation of peri-AIS astrocytes (pA30-pA80, t-test, P<0.01; P<0.05; **Fig. 4E**), whereas no changes were observed when the S100β antibody was applied prior to optical stimulation (all steps; t-test, P>0.05; **Fig. 4F**). Changes in rheobase values were observed in recorded neurons after optogenetic activation of peri-AIS astrocytes (33 ± 15 pA before vs. 25 ± 10 pA after stimulation, n=6, t-test, P<0.05; **Fig. 4E**), whereas no significant changes were detected following prior application of the S100β antibody before activation (**Fig. 4F**). It might be expected that S100β is not the only gliotransmitter released by astrocytes to regulate this form of LTP-IE and rapid AIS plasticity. Given that ATP/adenosine released by astrocytes can alter AIS excitability (Lezmy et al., 2021), we decided to test the local administration of Suramin (100 µM) (a purinergic antagonist) for 5 minutes before optogenetic stimulation of astrocytes. However, our results showed that the administration of Suramin did not abolish either LTP-IE or the morphological changes in the AIS (**Fig. S3**). These findings demonstrate that selective activation of peri-AIS astrocytes is sufficient to induce LTP-IE and AIS remodeling, highlighting the crucial involvement of astrocyte-derived S100β in regulating neuronal excitability and axonal plasticity.

### The astrocytic protein S100β rescues LTP-IE and rapid AIS plasticity

To further confirm the key role of S100β in LTP-IE and rapid AIS remodeling observed in V1 L5PNs, we repeated the previous experiment of astrocytic network inhibition but with local application of S100β (129 μM) for 5 minutes prior to the induction protocol. This approach allowed us to fully saturate the local environment without inducing spiking during the protocol itself, which could otherwise alter the results. Consistent with our previous findings, application of S100β to the proximal axon, while peri-AIS astrocytes were inactivated, was sufficient to rescue LTP-IE (157% ± 12, n=6 vs 102% ± 6, n=5, (MW) U test, P<0.01, **Fig. 5A**). Confocal imaging also revealed rescue of AIS remodeling with S100β application (mean AIS length: 33.45 ± 1.78 µm before induction and 36.57 ± 2.10 µm after induction, n=6, t-test, P<0.01; **Fig. 5B**). Input-output curves showed that responses to suprathreshold current amplitudes significantly increased after application of S100β (pA40-pA100, t-test, P<0.05; **Fig. 5C**). Additionally, significant differences were observed in rheobase values (45 ± 16 pA before vs. 31 ± 14 pA after induction, n=6, t-test, P<0.05; **Fig. 5D**). To insure that the effects of the locally applied S100β resulted from its Ca²⁺ chelating action, we confirmed these findings with the local extracellular application of BAPTA (5 mM) which exerts similar effects on Ca²⁺ (**Fig. S4**). Together, these results further support the role of S100β as a key regulator of LTP-IE and rapid AIS remodeling.

**Figure 5.**
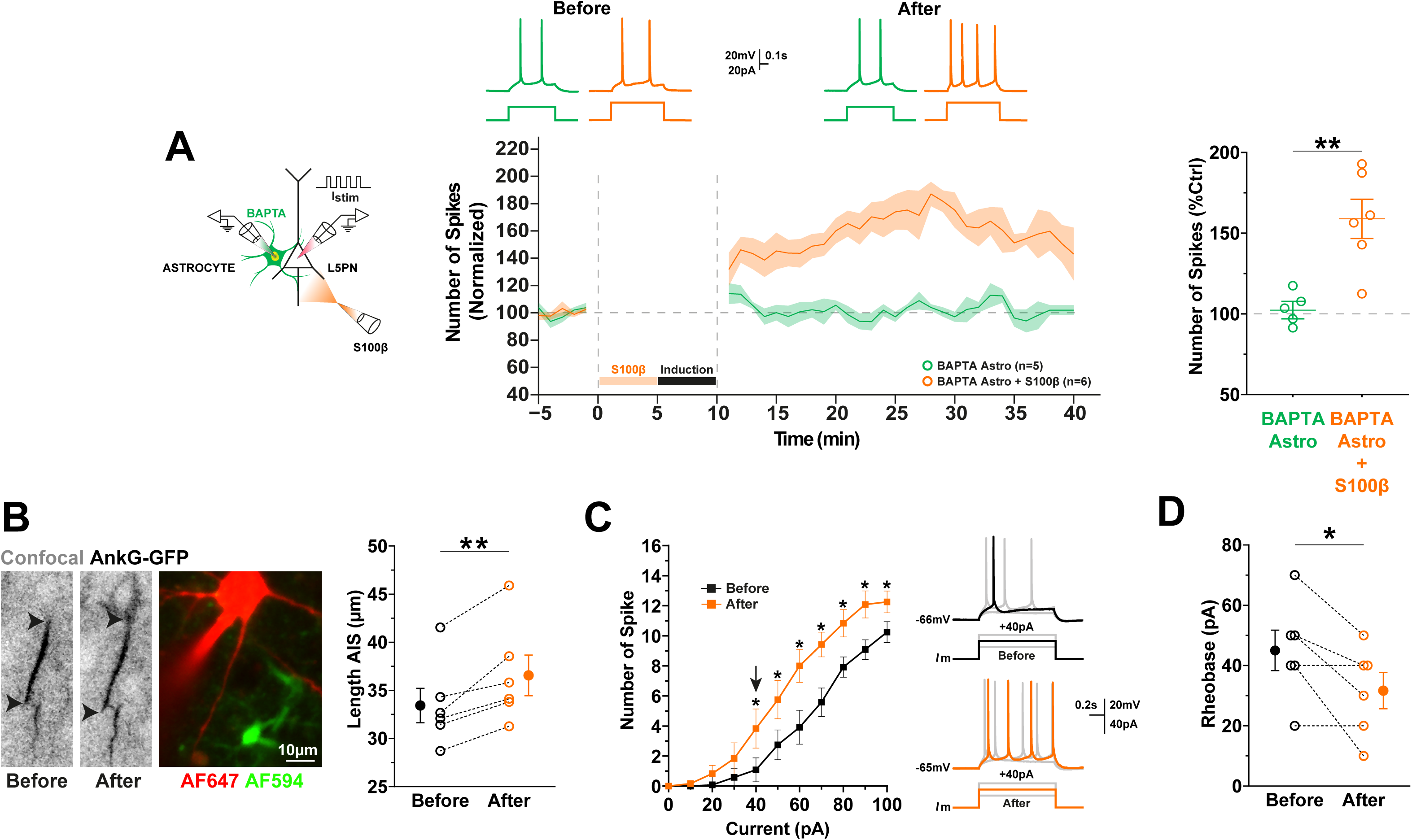
S100β rescues LTP-IE and AIS plasticity. **A.** *Left.* Experimental paradigm. Stimulation was performed by current injection into the whole-cell recorded neuron, with BAPTA-filled peri-AIS astrocyte inactivation and S100β protein applied 5 minutes before induction. *Middle.* Time course of normalized spike number before and after induction during peri-AIS astrocyte network inactivation (green) and following S100β application (orange), with representative traces shown above. *Right.* Quantification of spike number changes in the last 10 minutes, with individual values for each cell represented as green (astrocytes inactivated) and orange (astrocytes inactivated with prior S100β application) open circles. **B.** *Left*. Live image of an AnkG-GFP-positive neuron filled with Alexa Fluor 647 before and after induction during peri-AIS astrocyte (filled with Alexa Fluor 594) network inactivation, with prior S100β application. Black arrowheads indicate the start and end of the AIS. *Right.* S100β rescues the elongation of AIS length. **C.** *Left.* Current-number of spike relationships for each condition (before and after induction, during peri-AIS astrocyte network inactivation, with prior S100β application). Induction caused a leftward shift in the mean curve. *Right.* Representative traces of individual current injections at different amplitudes. Note the difference in the spike number with the same amplitude before and after induction, during peri-AIS astrocyte network inactivation, with prior S100β application. **D.** Change in Rheobase in the recorded neuron during peri-AIS astrocyte network inactivation with prior S100β application.

## DISCUSSION

Previous studies have shown that the AIS is a highly dynamic structure that contributes to the fine regulation of neuronal output (Grubb and Burrone, 2010; Evans et al., 2013). However, until now, plastic changes in the AIS have been limited to static post-hoc snapshots in large, differentially treated populations, preventing the detection of small-scale changes that may be obscured due to intercellular heterogeneity. Here, we have leveraged a new AnkG-GFP mouse model (Thome et al., 2025) to monitor the AIS in real time at the single-cell level. We found that in the visual cortex, the long-lasting increase of intrinsic excitability (LTP-IE) produced by a brief period of high frequency activity as previously described by Cudmore and Turrigiano (2004) is associated with rapid structural plasticity of the AIS, predominantly characterized by a distal elongation of its length (**Fig. 1**). This observation aligns with previous studies (Kuba et al., 2015; Jamann et al., 2021) showing that AIS elongation is linked to increased intrinsic excitability and a lower action potential (AP) threshold, highlighting a coordinated interplay between functional and structural plasticity of the AIS. Astrocytes have long been recognized as regulators of synaptic transmission, plasticity, and excitability through gliotransmitter release and local ion homeostasis. However, to our knowledge, their potential involvement in the plasticity of intrinsic excitability, including AIS morphological changes, has not yet been explored. We have demonstrated that peri-AIS astrocytes (proximal or adjacent to the AIS) can regulate the structural and functional plasticity of the AIS following brief periods of intrinsic spiking activity. This regulation depends on the release of the astrocytic calcium-binding protein S100β.

### LTP-IE is associated with increases AIS

Our results demonstrate that LTP-IE in L5PNs of the visual cortex is associated with a rapid increase in AIS length. This finding is significant, as it highlights the AIS’s ability to very rapidly (within minutes) adjust its length to regulate neuronal excitability, while it was believed to occur only on longer timescale (hours to days) acting mostly as a homeostatic mechanism. These results demonstrate that AIS morphology plasticity is also a mechanism allowing a neuron to adapt rapidly its excitability in addition to others such as regulation of ion channels to modulate spike thresholds or EPSP amplification (Debanne and Russier, 2019). LTP-IE in non-homeostatic form has been described in other brain regions including the hippocampus and the brainstem (Nelson et al., 2003; Sourdet et al., 2003) and it would be interesting to know if it is accompanied by changes in AIS morphology as well. The mechanisms described for LTP-IE so far seem to vary from a region to another involving Protein Kinase A in the visual cortex (Cudmore and Turrigiano, 2004), metabotropic glutamate receptor subtype 5 (mGluR5) in the somatosensory cortex, NMDA receptors in the hippocampus (Sourdet et al., 2003; Xu et al., 2005), down-regulation of Kv1 channels in the dorsal lateral geniculate nucleus (Duménieu et al., 2025) or reduction in BK-type calcium-activated potassium currents in the brainstem (Nelson et al., 2003). However, all these changes have ultimately been associated with the modulation (increase or decrease) of intracellular or extracellular calcium, as they are inhibited by calcium chelation. This could also implicate the astrocytic calcium-binding protein S100β, which is strongly expressed in all these plastic regions, and that we demonstrated is required for LTP-IE in the visual cortex. But S100β is much more than a simple Ca^2+^ chelator (Sorci et al., 2010).

### S100β: a multifaceted protein

Given that extracellular calcium is a key regulator of all forms of plasticity, S100β, which reduces [Ca^2+^]_e_, is expected to have a significant impact on these processes. Surprisingly, our results demonstrated that S100β does not inhibit LTP-IE or AIS plasticity in the visual cortex; rather, it is essential for these processes despite that both have been shown to depend on calcium elevation and Ca^2+^-activated molecules such as calcineurin and cyclin-dependent kinase 5 (cdk5) (Chand et al., 2015). This underscores the complex and multifaceted role of S100β, which extends beyond calcium regulation. S100β has been shown to interact directly or indirectly with various voltage gated channels, including Ca²⁺, K⁺, and Na⁺ channels (Hermann et al., 2012). For example, it facilitates Ca²⁺ flux across artificial membranes, activates L-type Ca²⁺ channels, and inhibits K⁺ channels such as the ether-à-go-go (hEAG1) and Kv4.2 channels (Bancroft et al., 2022). Our previous results demonstrated that S100β enhances the spiking duration and frequency of L5PNs through NaV1.6 potentiation (Ryczko et al., 2021; Gaudel et al., 2025). This is explained by the fact that Na^+^ channels are transiently clogged by Ca^2+^ ions occupying the channel lumen (Armstrong, 1999; Armstrong and Cota, 1999). Notably, LTP-IE requires neurons to fire at approximately 20–30 Hz during the induction protocol. The release of S100β could further promote high-frequency firing and facilitate LTP-IE induction. This idea is further supported by the fact that BAPTA rescues LTP-IE **(Fig S4)**. In addition, activity-dependent relocation of the AIS has been shown to be mediated by T- and L-type calcium channels (Grubb and Burrone, 2010). Similar to what has been described for S100A1, another member of the S100 protein family, S100β could directly bind to L-type calcium channels and increase their opening probability through protein kinase A (PKA) activation (Treves et al., 1997) which is required for LTP-IE activity in the visual cortex (Cudmore and Turrigiano, 2004). S100β secretion is tightly linked to neuronal activity (Nardin et al., 2009), and could serve as a gatekeeper of intrinsic plasticity to allow rapid AIS plasticity in response to high periods of activity, a processus which could be really important during development, especially in the visual cortex (Gutzmann et al., 2014). In addition, S100β could provide a direct mechanism for astrocytes to modulate synaptic plasticity through Ca^2+^ regulation. S100β has long been considered as a direct competitor of calmodulin, another Ca^2+-^binding protein that plays a key role in synaptic plasticity (Xia and Storm, 2005), which can tilt the balance toward LTD or LTP depending on the level of Ca^2+^ elevation. In line with this idea, S100β perfusion enhanced LTP in the hippocampus (Nishiyama et al., 2002). Taken together, these findings suggest that S100β may also influence other forms of plasticity beyond LTP-IE. Given the well-established interplay between synaptic and intrinsic plasticity (Debanne et al., 2019), it would be interesting to explore whether the interplay between synaptic and intrinsic plasticity involves structural modifications at the AIS, and whether S100β, or other gliotransmitters, act as key mediators of these coordinated forms of plasticity.

### S100β, NaV 1.6 and AIS reorganization

Na^+^ channels are transiently clogged by Ca^2+^ ions occupying the channel lumen. Reduction of extracellular Ca^2+^ relieves this partial block of the Na^+^ channel and allows for a larger influx of Na^+^ currents (Armstrong, 1999; Armstrong and Cota, 1999), and an enhanced sodium persistent current (I_NaP_), which is associated with the control of neuronal excitability and rhythmic activity (Agrawal et al., 2001; Wu et al., 2005). I_NaP_ is most commonly associated to Na^+^-channels including the Nav1.6 α-subunit (Drouillas et al., 2023), which is abundantly located at the distal end of the AIS (Van Wart et al., 2007; Hu et al., 2009; Drouillas et al., 2023), where the initiation of APs in L5PNs has been experimentally demonstrated (Palmer and Stuart, 2006), coinciding with the region where we observed structural changes in the AIS in our experiments (**Fig. 1C**). Furthermore, according to computational and experimental analyses, Nav1.6 α-subunits determine the lowest threshold for AP initiation in the distal AIS of cortical pyramidal neurons (Hu et al., 2009). We observed a reduction in the threshold potential for evoking APs (**Fig. 1E**), which could indicate an increase in the density and/or reinsertion of these channels after inducing LTP-IE. The absence of changes in passive cell properties (see Methods) further supports that LTP-IE associated with AIS elongation is mediated by changes in ionic conductances. Molecular trafficking of protein complexes in the AIS can follow two routes during development: (1) either insertion of proteins such as neurofascin 186 (Nfasc186) or the Kv7.3 channel into the somatodendritic and/or axonal membrane followed by bidirectional diffusion until immobilization by interaction with AnkyrinG (AnkG) (Ghosh et al., 2020); or (2) preferential vesicular insertion directly into the AIS, allowing binding to AnkG, as in the case of Nav1.6 (Akin et al., 2015). The maintenance of Nav1.6 channels has not yet been thoroughly studied, but so far, it appears that once anchored, they do not migrate to adjacent membranes. Therefore, although one study showed that prolonged activity caused lateral diffusion of other receptors (GABA_A_) in the AIS of hippocampal neurons (Muir and Kittler, 2014), we hypothesize that the origin of these structural changes was rather a vesicular insertion of new Nav1.6 channels into the distal AIS minutes after the period of elevated activity. This is consistent with the results of Kuba et al., (2010), who showed that AIS elongation was accompanied by increased expression of Na^+^ channels and enhanced axonal I_NaP_ and membrane excitability in the auditory pathway. Moreover, a computational model predicted that the distal AIS exhibits greater variation in terms of Na^+^ conductance and that small changes in the density of functional Na^+^ channels in this region would have a strong impact on neuronal excitability in L5PNs (Baranauskas et al., 2013). We do not rule out the simultaneous contribution of other VGCCs, such as the combination of T-type and/or L-type VGCCs, which were necessary for activity-dependent AIS relocation (Grubb and Burrone, 2010; Kuba et al., 2015), or K^+^ channels, such as Kv7.2, whose activity-dependent expression can increase in the AIS, reducing the shunt current around the AP threshold (Jamann et al., 2021) and enhancing the electrotonic isolation of the distal AIS, a factor that had also been computationally predicted as a key modulator of neuronal excitability (Baranauskas et al., 2013), or Kv2.1, whose phosphorylation, localization in the AIS (non-canonical pathway), and function are regulated by neuronal activity and can drastically affect excitability and intrinsic plasticity (Misonou et al., 2005; Jensen et al., 2017). Even other interactions between key protein partners could have contributed to this activity-driven intrinsic plasticity, such as the phosphorylation of casein kinase 2 (CK2), which regulates the interaction between Na^+^ channels and AnkG (Bréchet et al., 2008), or the interaction of Nav1.6 with the microtubule-associated protein Map1b (Mtap1b), which facilitates the trafficking of these channels to the cell surface (O’Brien et al., 2012; Solé et al., 2019). Leveraging the opportunities offered by novel genome-editing techniques and super-resolution microscopy (Willems et al., 2020; Liu et al., 2022), future studies should investigate the exocytic machinery in the AIS in contexts of activity-dependent intrinsic plasticity, perhaps starting with endogenous tagging of Nav1.6 in L5PNs.

### AIS plasticity in critical periods

Brain plasticity driven by activity or experience is enhanced during critical periods. For over 40 years, the primary model for studying critical period plasticity has been the primary visual cortex (Wiesel and Hubel, 1963; Hensch, 2005). Specifically, the critical visual period in mice has been defined from eye opening (P13) to approximately P35. The P13-P19 phase is considered a precritical period, independent of visual experience; P19-P28 represents the critical period with the highest sensitivity to imbalances; and P28-P35 corresponds to the consolidation and closure of the critical period (Gordon and Stryker, 1996; Corlew et al., 2007). Interestingly, the most significant elongation and reduction in AIS development in the visual cortex occur approximately one week before the opening and closing of the critical period, respectively (Gutzmann et al., 2014). This structural refinement of the AIS may be necessary to prepare the cortical network for the subsequent initiation and termination of the critical period. However, what determines the duration of these plasticity windows? GABAergic inhibition has gathered substantial evidence as a key mechanism regulating the onset and duration of the critical period in the visual cortex (Hensch, 2004, 2005). However, it is not the only factor. A recent study demonstrated that astrocytic connexin 30 controls the timing of critical period closure in the visual cortex (Ribot et al., 2021), and remarkably, the reintroduction of immature cultured astrocytes reinstates a period of heightened plasticity in adult animals, resembling the critical visual period (Müller and Best, 1989). Moreover, in other brain regions such as the hippocampus and somatosensory cortex, astrocytes have also been implicated in the closure of the critical plasticity period (Falcón-Moya et al., 2020; Martínez-Gallego et al., 2022). While we have not explored the role of astrocytes in AIS plasticity beyond the visual critical period, it would be interesting to investigate whether astrocytic manipulation could reinstate a period of AIS plasticity in adulthood, when its length is presumed to be stable.

## CONCLUSION

The million-dollar question is how astrocytes detect neuronal activity and, more importantly, how they determine when to release the appropriate gliotransmitters, specifically, S100β in this case, at the right time. As previously mentioned, astrocytes can detect neuronal activity through changes in the extracellular ion composition, including K⁺, Ca²⁺, and Na⁺. In particular, the sodium/calcium exchanger (NCX) is predominantly expressed in the distal processes and terminal feet of astrocytes and can be easily activated by an increase in astrocytic [Na^+^]i driven by neuronal activity. Given that certain astrocytic processes can contact the AIS, it is tempting to speculate that intense neuronal activity increases [Na^+^]_i_ signals in astrocytes, allowing Ca^2+^ influx from the extracellular space through the NCX exchanger. In future experiments, it would be interesting to determine whether [Na^+^]_i_ and [Ca^2+^]_i_ transients in astrocytes are closely related to specific protocols of intrinsic plasticity.

## MATERIAL AND METHODS

### Experimental model

All experiments were conducted in accordance with the rules of the Canadian Institutes of Health Research and were approved by the Animal Care and Use Committee of the University of Montreal (Protocol #23-207). We crossed mice from the newly generated AnkG-GFP-lox knock-in line (B6;129SV-ank3tm1DCI/HD) (Thome et al., 2025) with Vglut2-cre mice (B6J.129S6(FVB)-Slc17a6tm2(cre)Lowl/Mwar). This allowed Cre-dependent expression of AnkG-GFP driven by the VGlut2 promoter, a genetic marker of glutamatergic signaling. VGlut2-AnkG-GFP were fed ad libitum and housed with a 12 h light/dark cycle. Male and female mice were used for experiments.

### V1 slice preparation

Coronal slices (350 µm) containing layer 5 of the V1 were prepared from 14-to 30-day-old Vglut2-AnkG-GFP mice. Mice were anesthetized with isoflurane (Pharmaceutical Partners of Canada Inc., Richmond Hill, ON, Canada) and subsequently decapitated. Following decapitation, the skull was quickly removed, and the brain was transferred to modified artificial cerebrospinal fluid (aCSF, in mM: 3 KCl, 1.25 KH2PO4, 4 MgSO4, 26 NaHCO3, 10 dextrose, 0.2 CaCl2, 219 sucrose; pH 7.3-7.4, 300-320 mOsmol/kg) that was ice-cold (0-4°C) and oxygenated with a mixture of 95% O2 and 5% CO2. Brain slices were then sectioned using a VT1000S vibratome (Leica). After a 15-minute incubation at 30°C in a temperature-controlled water bath within a slice holder containing oxygenated artificial cerebrospinal fluid (aCSF, in mM: 124 NaCl, 3 KCl, 1.25 KH2PO4, 1.3 MgSO4, 26 NaHCO3, 10 dextrose, and 1.2 CaCl2; pH 7.3-7.4, 294-300 mOsmol/kg) bubbled with 95% O2 and 5% CO2, slices were transferred to the recording chamber at room temperature. A recovery period of at least one hour was allowed for the slices before recording.

### Electrophysiology

Recordings were conducted at room temperature in a submerged chamber continuously perfused with artificial CSF (in mM: 124 NaCl, 3 KCl, 1.25 KH2PO4, 1.3 MgSO4, 26 NaHCO3, 10 dextrose, and 1.2 CaCl2; pH 7.3-7.4, 294-300 mOsmol/kg) bubbled with 95% O2 and 5% CO2. Patch microelectrodes were fabricated from borosilicate glass capillaries (1.5 mm outside diameter, 1.12 mm inside diameter, World Precision Instruments) using a P-97 puller (Sutter Instruments). For neuronal recordings, pipettes with resistance ranging from 6 to 12 MΩ were filled with an internal solution containing the following (in mM): 140 K-gluconate, 5 NaCl, 2 MgCl2, 10 HEPES, 0.5 EGTA, 2 Tris ATP salt, 0.4 Tris GTP salt, pH 7.2-7.3, 280-300mOsmol/kg. To visualize neuronal and axonal morphologies during the experiment, Alexa Fluor 647 was added to the internal solution. Layer 5 pyramidal neurons of the extratelencephalic subtype were selected based on their morphological characteristics (large soma and thick apical dendrite) and electrophysiological properties (burst spiking). All recordings were performed using a Multiclamp 700A amplifier, Digidata 1550B interface coupled to a computer equipped with pClamp 11 software (Molecular Devices, San Jose, CA). After the establishment of a gigaseal, the membrane potential was held at -65 mV, and the membrane patch was ruptured by suction. During the experiments, the changes in Vm (1–6 mV) were corrected by imposing continuous current to maintain the membrane potential constant at -65mV. Recordings were discarded if the resting membrane potential shifted by >6 mV or if resting input resistance (measured with a 50pA hyperpolarizing pulse lasting 50 ms, delivered every 10 seconds) varied >20%. Astrocytes were initially identified morphologically after incubating the slices in the marker Sulforhodamine 101 (SR101) for 5 minutes, based on their size and shape. This identification was subsequently confirmed by their electrophysiological characteristics, including a low resting membrane potential and passive responses to negative and positive current injections. For experiments involving astrocyte inactivation, BAPTA tetrapotassium salt (20 mM) was added to the internal solution filling the recording pipette. All recordings were performed in the presence of antagonists of N-methyl-d-aspartate (NMDA), AMPA/kainate, and GABA_A_ receptors [2-amino-5-phosphonovaleric acid (d-APV), 50 μM; 6-cyano-7-nitroquinoxalene-2,3-dione (CNQX), 20 μM; and Gabazine, 20 μM, respectively, which were all purchased from Tocris Biosciences (Ellisville, Missouri, USA)].

### Plasticity Protocol and Analysis

After achieving whole-cell access by rupturing the membrane with negative pressure, a waiting period of 5 to 10 minutes was allowed for the cell to dialyze with the pipette solution and for the Alexa 647 fluorescent dye to diffuse. Throughout the recording, intrinsic excitability was assessed every 10 seconds using a small, constant-amplitude depolarizing pulse (500 ms, 10–60 pA), which was selected to evoke 2–3 APs during the baseline period (5 minutes) and then maintained constant throughout the recording. To generate input-output curves and determine the current threshold for AP generation, a series of depolarizing current injections (0–100 pA in 10 pA increments) was applied before the baseline period and at the end of the recording. Each series was delivered at least twice per cell, and responses were averaged. The induction stimulus consisted of 500 ms depolarizing pulses (150–180 pA) delivered 60 times at 4-second intervals, with an amplitude selected to evoke a sustained firing rate of 18–30 Hz. Only neurons with a stable baseline, consistent recording parameters, and a post-induction recording duration of at least 20 minutes were included in the analysis. The degree of LTP-IE was calculated as the ratio of firing frequency during the last 10 minutes of the post-induction recording to the baseline. Intra-and intercellular comparisons were performed by extracting the average response per minute (6 pulses every 15 seconds), and the mean firing rate during the 5-minute baseline period was used to normalize data before and after induction. Control cells remained at rest during the induction stimulus period, and analyses were conducted similarly in this condition.

### Optogenetic Stimulation

Astrocytes peri-AIS in VGlut2-AnkG-GFP mice, which received bilateral intracerebral viral administration at P0 of an AAV2/5-GFAP-hChR2(H134R)-mCherry (3.40E+12) expressing GFAP-promoter-driven Channelrhodopsin-2 (ChR2) fused to the mCherry reporter (provided by the CNP Viral Vector Core at the CERVO Research Center, RRID: SCR_016477), were optogenetically stimulated (beginning two weeks after viral injection) using two lasers (440 nm and 488 nm) simultaneously in the SIM light path of an FV1000 microscope (Olympus). The SIM scanner was used in square scanning mode to photoactivate small manually delineated areas within astrocytes near the AIS of the recorded neurons. Optogenetic stimulations were applied after the baseline period using 60 pulses of 2 s ON and 2 s OFF (10–20% laser power / 8.6–14.9 µW for the 440 nm laser / 8.7–15.9 µW for the 488 nm laser). To assess the efficacy of the stimulation, laser pulses were applied while recording the electrophysiological activity of astrocytes. These recordings confirmed that the photostimulation reliably evoked small membrane depolarizations, averaging 7.4 mV and ranging between 6 and 9 mV.

### Drugs

Chemicals used in this study were purchased from Sigma-Aldrich (Oakville, Ontario, Canada), Tocris Biosciences (Ellisville, Missouri, USA) and Abcam (Cambridge, UK). The following drugs were bath-applied using a syringe pump: D,L-2-amino-5-phosphonovaleric acid (APV, 75 µM), CNQX (10 µM), SR 95531 hydrobromide (Gabazine, 20 µM) and Suramin (100 µM). In some experiments, the Ca²⁺-binding protein S100β (129 μM) (Produced in the Department of Biochemistry of the Université de Montreal as described in Morquette et al., 2015) or 1,2-bis(o-aminophenoxy) ethane-N,N,N′,N′-tetraacetic acid tetrasodium salt (BAPTA, 5 mM) were locally applied using glass micropipettes (tip diameter ∼1 μm) with 2–20 psi pressure pulses for 5 minutes (Picospritzer III, Parker Instrumentation, Fairfield, NJ, USA). Monoclonal anti-S100β antibodies (mouse anti-S100β, Sigma Aldrich #S2532 or rabbit anti-S100β, Abcam ab56642), as well as their denatured form, were applied locally with large-tip (10-20 µm) glass micropipettes carefully lowered near the AIS recorded neuron with 0.1-2 psi pressure pulses lasting 5 minutes. All bath-applied chemicals were diluted in water at 100X their final concentrations and further diluted through the perfusing aCSF at their final concentrations.

### Immunohistochemistry and confocal microscopy

For the immunohistochemistry against AnkG and NeuN, coronal brainstem slices (350 μm) were prepared from 14 to 30-day-old VGlut2-AnkG-GFP mice using a vibratome VT 1000S (Leica), immediately immersed in a solution of 4% (wt/vol) paraformaldehyde in PBS, and kept for one hour at 4°C. The sections were rinsed 3 times for 10 minutes in PBS and incubated in a blocking solution containing 0.3% Triton X-100 and 5% Normal Donkey Serum (Jackson ImmunoResearch #017-000-121) in PBS for one hour at room temperature. The sections were then rinsed 3 times for 10 minutes in PBS and incubated overnight at 4°C in a mix of the primary antibodies (chicken anti-AnkG, EnCor Biotech #PCAC-ANK3, dilution 1:1000; mouse anti-NeuN, Millipore #MAB377, dilution 1:500). The following day, the slices were rinsed 5 times for 10 minutes in PBS and incubated in the relevant secondary antibodies mix (donkey anti-chicken Alexa Fluor 647, Jackson Immunoresearch #703-605-155, dilution 1:250; donkey anti-mouse Alexa Fluor 594, Jackson Immunoresearch #715-585-151, dilution 1:250) diluted in the blocking solution for 120 minutes in a dark chamber at room temperature. The sections were then rinsed 4 times for 15 minutes in PBS and mounted on ColorFrost Plus slides (Fisher Scientific, Ottawa, Ontario, Canada) using Fluoromount-G (Southern Biotech, Birmingham, Alabama, USA). A similar procedure was used for the immunohistochemistry against S100β, except that the slices were immersed in 4% (wt/vol) paraformaldehyde in PBS overnight at 4°C. The staining was performed using mouse anti-S100β (Sigma #S2532, 1:400) as the primary antibody and donkey anti-mouse Alexa Fluor 647 (Abcam #ab150107, 1:250) as the secondary antibody. The washing steps, blocking conditions, and mounting procedure were identical to those described above. Slides imaging was performed using either a Zeiss LSM-900 Confocal Airyscan microscope or an FV1000 confocal microscope (Olympus). AnkG-GFP was visualized in real-time during electrophysiology experiments using the FV1000, operated with FV10-ASW imaging software (Version 03.00). Z-stacks were acquired at a speed of 10 μs/pixel and a resolution of 800 × 800 pixels, with 1 μm step sizes and a zoom range of 2 to 4×. To minimize phototoxicity and photobleaching, AIS z-stacks were captured at two time points: 5 minutes before induction or drug exposure, as applicable, and 30 minutes afterward. AIS length was measured manually and by fluorescence intensity along a 1-pixel-wide line drawn from the soma down the axon (1 pixel ≈ 0.206 μm), using Fiji software (ImageJ 1.50e, NIH, USA). Fluorescence profiles were smoothed using a 10-point (∼2 μm) sliding mean and normalized between 1 (maximum) and 0 (minimum). The AIS end was defined at a 33% fluorescence threshold (method adapted from Grubb and Burrone, 2010). No difference was observed between the two methods. In a subset of experiments (**Fig. S1**), images were acquired using a two-photon microscope equipped with a polygon scanner (BliQ Photonics), enabling high-speed imaging while minimizing phototoxicity from repeated AIS imaging during recording.

### Quantification and statical analysis

Data are presented as mean ± standard error to the mean (SEM) and as proportions (%). Statistical analyses were performed depending on the experimental design. All paired comparisons before and after treatment were confirmed to follow a normal distribution (Shapiro–Wilk test, P>0.05) and were analyzed using paired t-tests. Comparisons between independent groups, for which normality could not be assumed, were performed using the non-parametric Mann–Whitney U test. Statistical significance was set at *P<0.05, **P<0.01, and ***P<0.001; NS indicates not significant. Data analysis was performed using SPSS (IBM SPSS Statistics).

## DECLARATIONS

### Author contributions

R.S.G., Y.I and A.K wrote the paper. R.S.G and Y.I collected and analyzed the data and built the figures. All authors contributed to the article and approved the submitted version.

## Acknowledgements

We thank Dr. Dorly Verdier for critically reading a preliminary version of the manuscript and members of A.K laboratories for comments on the manuscript and technical support. We are especially grateful to our esteemed colleagues, Dr. James G. Omichinski and Dr. Haytham Wahba, from the Department of Biochemistry at the Université de Montréal, for their invaluable contribution to the production of the S100β protein, and to Dr. Paul Jenkins, from Michigan Neuroscience Institute Affiliate, for generously providing the AnkG-GFP-lox mouse line. The graphical abstract incorporates elements from SciDraw (Federico Claudi: https://doi.org/10.5281/zenodo.3925905; Andrew Hardaway: 10.5281/zenodo.5348394; Roberta Schellino: 10.5281/zenodo.10390020), an open-access platform for scientific illustrations.

## Funding

This work was supported by the Canadian Institutes of Health Research (Grant # 197855), Natural Sciences and Engineering Research Council of Canada (RGPIN/05255-2020), CIRCA Post-doctoral fellowship to Y.I.

## Conflict of Interest Statement

The authors declare that the research was conducted in the absence of any commercial or financial relationships that could be construed as a potential conflict of interest.

## Availability of data and materials

All data generated or analysed during this study are included in this published article.

## Preprint

This manuscript was posted on a preprint.

---

**Supplementary Figure 1.**
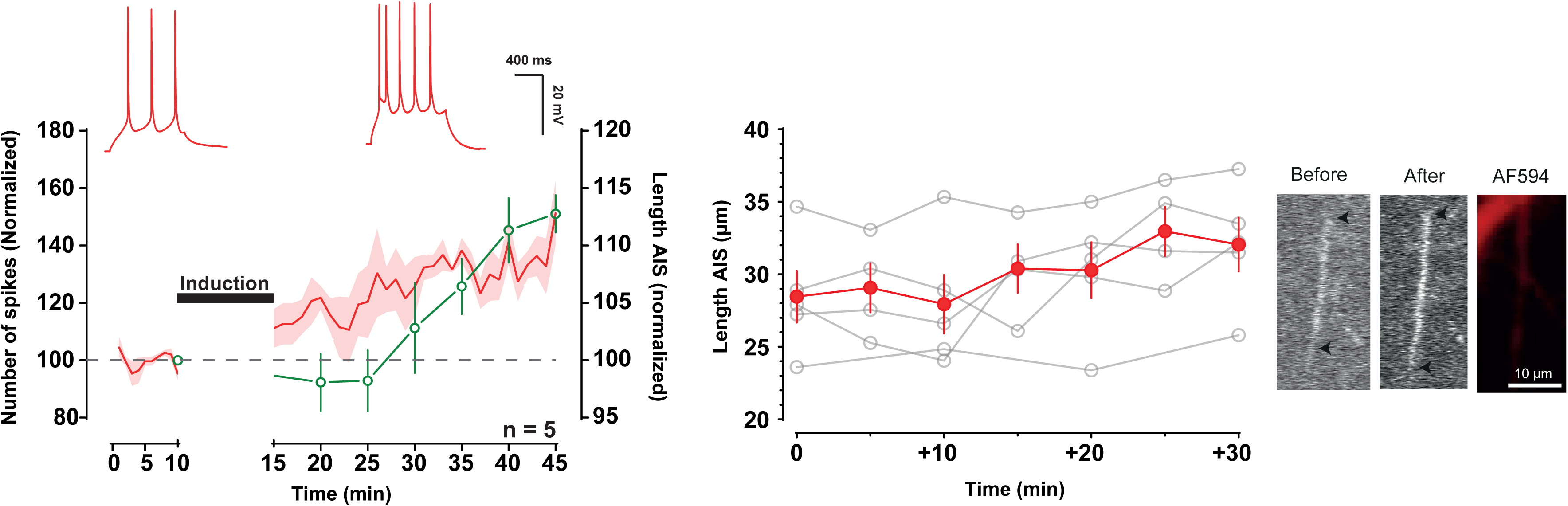
AIS plasticity linked to LTP-IE emerges within minutes post-induction. *Left.* Time course of normalized spike count before and after induction (red), with representative traces shown above. Time course of normalized AIS length (green) measured every 5 minutes post-induction. Note the immediate change in excitability following induction and the increase in AIS length occurring 10–15 minutes later. *Right.* Individual AIS length measurements over time, with light gray representing individual cells and red indicating the average. Example images are shown at baseline (0 min, before) and 30 minutes (after) post-induction.

**Supplementary Figure 2.**
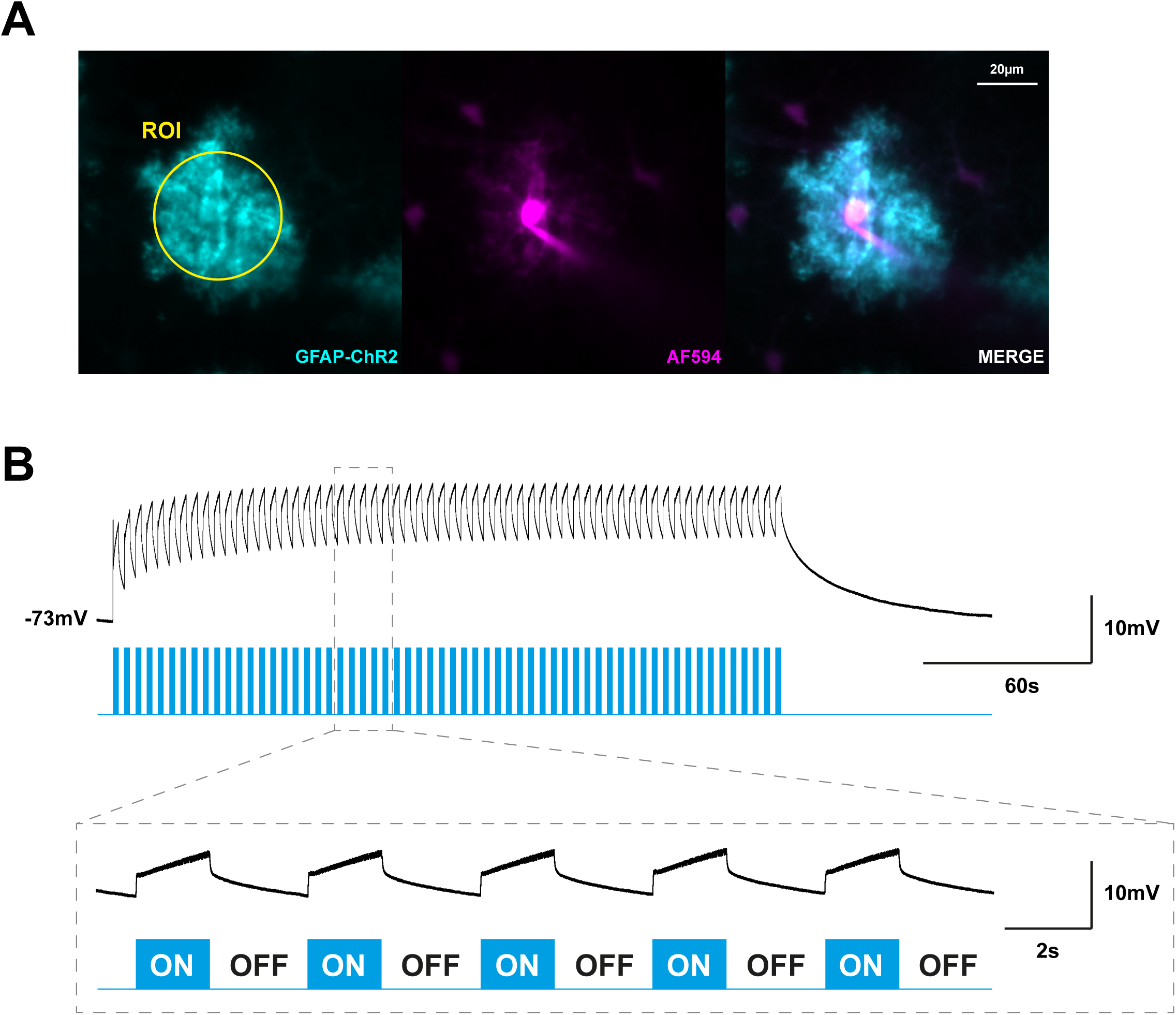
Representation of the astrocyte optogenetic stimulation protocol. **A.** Photomicrograph of a patched astrocyte in whole-cell configuration, filled with Alexa Fluor 594 and expressing ChR2 (cyan) for the data presented in **B**. The ROI drawing represents the optogenetic stimulation area. **B.** *Top*. Membrane responses of the astrocyte in **A** to optogenetic stimulation (ROI), representing the complete stimulation protocol used in our experiments with VGlut2-AnkG-GFP mice injected with AAV GFAP-ChR2. *Bottom.* Expansion of a part of the stimulation protocol clearly showing how each scanning episode (2s) triggers a membrane depolarization.

**Supplementary Figure 3.**
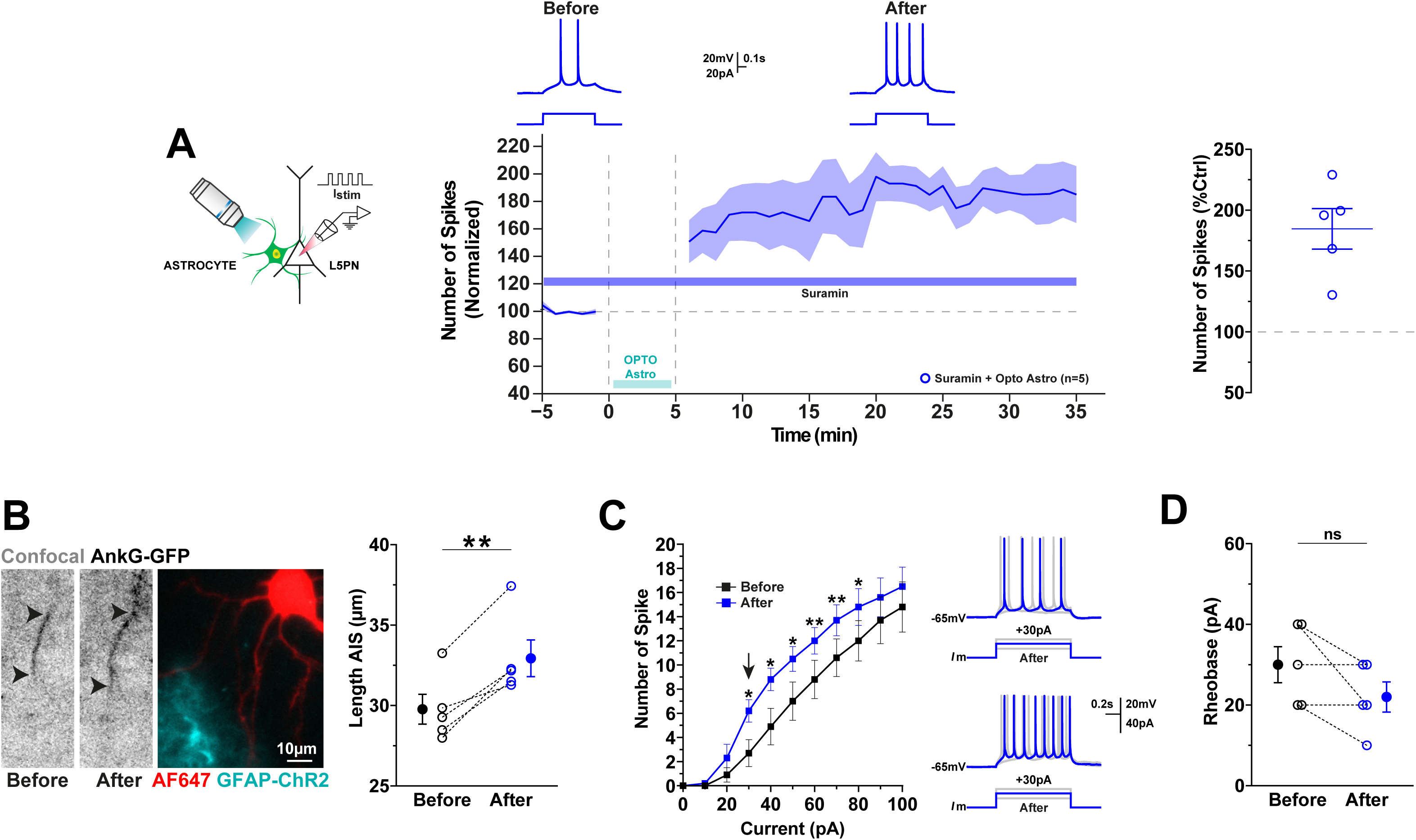
ATP/adenosine inhibition does not affect LTP-IE or AIS plasticity. **A.** *Left.* Experimental paradigm. Stimulation was performed by optogenetic activation of an astrocyte near the AIS of the recorded whole-cell neuron, after a 5-minute application of the ATP/adenosine antagonist Suramin. *Middle.* Time course of normalized spike number before and after induction during peri-AIS astrocyte optogenetic activation in the presence of Suramin (blue), with representative traces shown above. *Right.* Quantification of spike number changes in the last 10 minutes, with individual values for each cell represented as open blue circles (optogenetic activation of astrocytes in the presence of Suramin). **B.** *Left*. Live image of an AnkG-GFP-positive neuron filled with Alexa Fluor 647 before and after optogenetic activation of an astrocyte expressing ChR2 (cyan), with Suramin present. Black arrowheads indicate the start and end of the AIS. *Right:* Suramin does not affect the elongation of AIS length. **C.** *Left.* Current-number of spike relationships for each condition (before and after optogenetic activation of astrocytes, under Suramin exposure). Optogenetic activation caused a leftward shift in the mean curve. *Right.* Representative traces of individual current injections at different amplitudes. Note the difference in the spike number with the same amplitude before and after optogenetic activation of astrocytes, while Suramin was applied. **D.** No change in Rheobase in the recorded neuron during optogenetic activation of peri-AIS astrocyte network in the presence of Suramin.

**Supplementary Figure 4.**
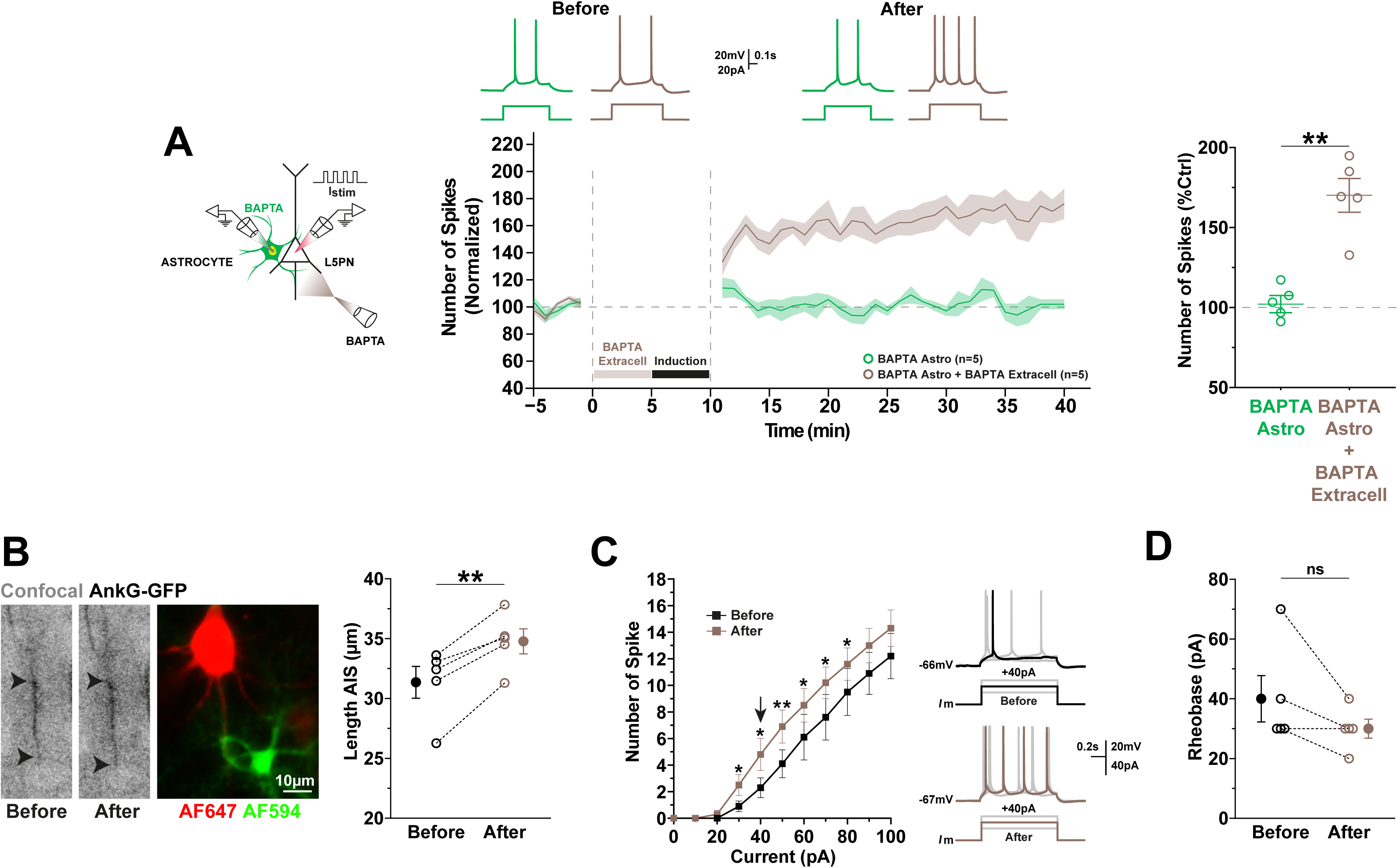
BAPTA extracellular rescues LTP-IE and AIS plasticity. **A.** *Left.* Experimental paradigm. Stimulation was performed by current injection into the whole-cell recorded neuron, with BAPTA-filled peri-AIS astrocyte for inactivation of the astrocytic syncytium and local extracellular application of BAPTA to mimic the Ca2+-chelating action of S100β, 5 minutes before induction. *Middle.* Time course of normalized spike number before and after induction during peri-AIS astrocyte network inactivation (green) and following extracellular BAPTA application (brown), with representative traces shown above. *Right.* Quantification of spike number changes in the last 10 minutes, with individual values for each cell represented as green (astrocytes inactivated) and brown (astrocytes inactivated with prior BAPTA extracellular application) open circles. **B.** Live image of an AnkG-GFP-positive neuron filled with Alexa Fluor 647 before and after induction during peri-AIS astrocyte (filled with Alexa Fluor 594) network inactivation, with prior extracellular BAPTA application. Black arrowheads indicate the start and end of the AIS. *Right.* BAPTA extracellular rescues the elongation of AIS length. **C.** *Left.* Current-number of spike relationships for each condition (before and after induction, during peri-AIS astrocyte network inactivation, with prior extracellular BAPTA application). Induction caused a leftward shift in the mean curve. *Right.* Representative traces of individual current injections at different amplitudes. Note the difference in the spike number with the same amplitude before and after induction, with prior BAPTA extracellular application. **D.** No change in Rheobase in the recorded neuron during peri-AIS astrocyte network inactivation with prior BAPTA extracellular application.

## Notes

### Competing Interest Statement

The authors have declared no competing interest.

### Summary of Updates

The manuscript title has been updated to better reflect the aim and objectives of the paper

